# Detailed analyses of stall force generation in *Mycoplasma mobile* gliding

**DOI:** 10.1101/142133

**Authors:** Masaki Mizutani, Isil Tulum, Yoshiaki Kinosita, Takayuki Nishizaka, Makoto Miyata

## Abstract

*Mycoplasma mobile* is a bacterium that uses a unique mechanism to glide on solid surfaces at a velocity of up to 4.5 µm/s. Its gliding machinery comprises hundreds of units that generate the force for gliding based on the energy derived from ATP; the units catch and pull on sialylated oligosaccharides fixed to solid surfaces. In the present study, we measured the stall force of wild-type and mutant strains of *M. mobile* carrying a bead manipulated using optical tweezers. The strains that had been enhanced for binding exhibited weaker stall forces than the wild-type strain, indicating that stall force is related to force generation rather than to binding. The stall force of the wild-type strain decreased linearly from 113 to 19 pN following the addition of 0–0.5 mM free sialyllactose (a sialylated oligosaccharide), with a decrease in the number of working units. Following the addition of 0.5 mM sialyllactose, the cells carrying a bead loaded using optical tweezers exhibited stepwise movements with force increments. The force increments ranged from 1 to 2 pN. Considering the 70-nm step size, this small unit force may be explained by the large gear ratio involved in the *M. mobile* gliding machinery.

**SIGNIFICANCE:** *Mycoplasma* is a genus of bacteria that parasitizes animals. Dozens of *Mycoplasma* species glide over the tissues of their hosts during infection. The gliding machinery of *Mycoplasma mobile*, the fastest species, includes intracellular motors and hundreds of legs on the cell surface. In the present study, we precisely measured force generation using a highly focused laser beam arrangement (referred to as optical tweezers) under various conditions. The measurements obtained in this study suggest that the rapid gliding exhibited by *M. mobile* arises from the large gear ratio of its gliding machinery.

## INTRODUCTION

Members of the bacterial class *Mollicutes*, which includes the genus *Mycoplasma*, are parasitic and occasionally commensal bacteria that are characterized by small cells and genomes, and by the absence of a peptidoglycan layer (1, 2). Dozens of parasitic *Mycoplasma* species, such as the fish pathogen *Mycoplasma mobile* (3–5) and the human pathogen *Mycoplasma pneumoniae* (6–8), have protrusions, and exhibit gliding motility in the direction of the protrusions on solid surfaces, which enables mycoplasmas to parasitize other organisms. Interestingly, *Mycoplasma* gliding does not involve flagella or pili, and is completely unrelated to other bacterial motility systems, or the conventional motor proteins that are common in eukaryotic motility. *M. mobile*, which can be isolated from the gills of freshwater fish, is a fast-gliding *Mycoplasma* (9–13). It glides smoothly and continuously on glass at an average speed of 2.0–4.5 µm/s, or 3–7 times the length of the cell per second. A working model, called the centipede or power-stroke model, has been proposed to explain the gliding mechanism of *M. mobile*: the cells repeatedly catch, pull, drag, and release the sialylated oligosaccharides (SOs) on solid surfaces (3–5, 14). The gliding machinery comprises internal and surface structures (15–19). The internal structure includes the α-and β-subunit paralogs of F-type ATPase/synthase, and generates the force for gliding, based on the energy derived from ATP (18–22). The force is probably transmitted across the cell membrane to the surface structure, which is composed of at least three huge proteins, Gli123, Gli521, and Gli349 (15–17, 23). Gli521, the crank protein, transmits the interior force of the cell to Gli349, with actual structural changes (17, 24). Gli349, the leg protein, extends after thermal fluctuation, and catches the SOs, which are the predominant structures on animal cell surfaces (Fig 1*A*) (9, 25–27). The cells always glide in the direction of the machinery, which may be a result of the directed binding of the cells on solid surfaces (3, 28). In theory, hundreds of gliding units on the cell surface should act cooperatively to enable smooth gliding (Fig. 1*B*) (15, 27, 29). To examine this working model in detail, it is necessary to investigate the behavior of the individual units. Recently, the discrete movements involved in gliding motility, which are possibly attributable to single leg movements, have been observed by controlling the working leg number following the addition of free SOs (22, 27). In the current study, we focused on the pulling force exerted on the solid surface, because the generation of force has not been properly investigated, with the exception of the final stall force of the wild-type strain (11). Therefore, in the present study, we quantitatively measured the pulling forces of several strains of whole cells under various conditions using optical tweezers (11, 28, 30–32), and characterized the force generated by the gliding machinery.

**Fig 1.**
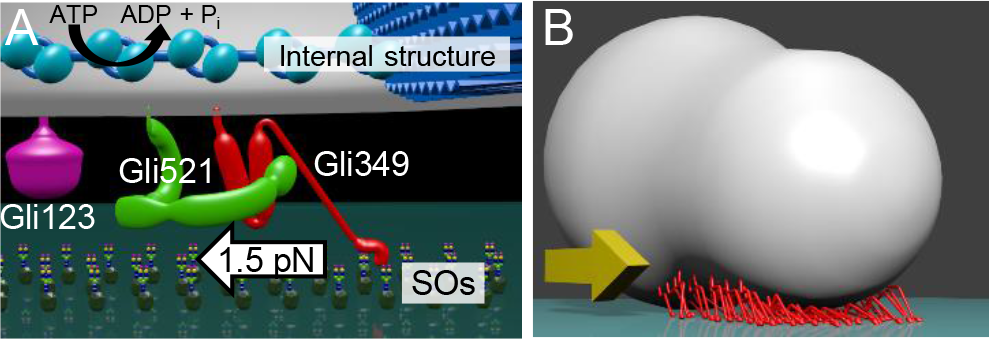
Schematic illustration of the gliding machinery. (*A*) Magnified illustration of a unit. The single unit consists of an internal structure (upper blue) and three huge proteins: Gli123 (purple), Gli349 (red), and Gli521 (green) on the cell surface. The force generated by the internal structure based on ATP hydrolysis is transmitted through Gli521 and pulls Gli349 repeatedly. Gli349 catches and pulls SOs on the solid surface; the unit force was estimated to be approximately 1.5 pN in the present study (see Fig. 6). (*B*) Approximately 75 legs (red) projecting from the cell can work simultaneously. The cell glides in the direction of the yellow arrow. Unbound legs are not illustrated.

## RESULTS

### Stall force measurement

The propelling force in *M. mobile* cells has been measured using optical tweezers (11) A bead bound to a cell was trapped by a highly focused laser beam, and the force was calculated by measuring the distance between the center of the bead and the trap; the force acting on the bead increased linearly with the displacement from the trap center (11, 28, 32). In the present study, *M. mobile* cells, which had been biotinylated and suspended in phosphate-buffered saline with glucose (PBS/G), were inserted into a tunnel chamber. Polystyrene beads coated with avidin (1-µm in diameter) were subsequently added to the tunnel. A bead trapped using optical tweezers was attached to the back end of the gliding cell by exploiting the avidin–biotin interaction (28, 33, 34). The cell pulled the bead from the trap center (Fig. 2*A* and Movie S1). Starting from 0 s, the pulling force increased and reached a plateau at approximately 40 s. The maximum value of the average of 25 data points was used as the stall force (Fig. 2*B*). The stall force of the wild-type strain was 113 ± 32 pN (*n* = 50).

To determine the proteins involved in force generation or force transmission, we measured the stall force in the wild-type strain of *M. mobile* to compare it with that in six previously isolated strains with known gliding speeds and/or binding activities (12, 21, 35). The *gli521* (P476R) mutant of protein Gli521 has a single amino acid substitution (proline for arginine at the 476th position) (21), but mutations in other strains have not been described. The pulling forces of the mutants increased from 0 s and stalled at 20–50 s. The stall force of the m14 strain—110 ± 29 pN (*n* = 29)—was not significantly different from that of the wild-type strain (*P* = 0.1 > 0.05, according to the Student’s *t*-test). The stall forces of the *gli521* (P476R), m6, m27, m29, and m34 mutants were significantly reduced to 64–81% of that of the wild-type strain (*P* = 6 × 10^-3^, 4 × 10^-4^, 9 × 10^-5^, 8 × 10^-9^, and 4 × 10^-4^, respectively; *P* < 0.01, according to the Student’s *t*-test) (Fig. 2*B Inset*). The *gli521* (P476R) mutant, which was characterized by enhanced binding, exhibited a smaller force than the wild-type strain, suggesting that the stability of binding is not the only determinant of stall force.

**Fig 2.**
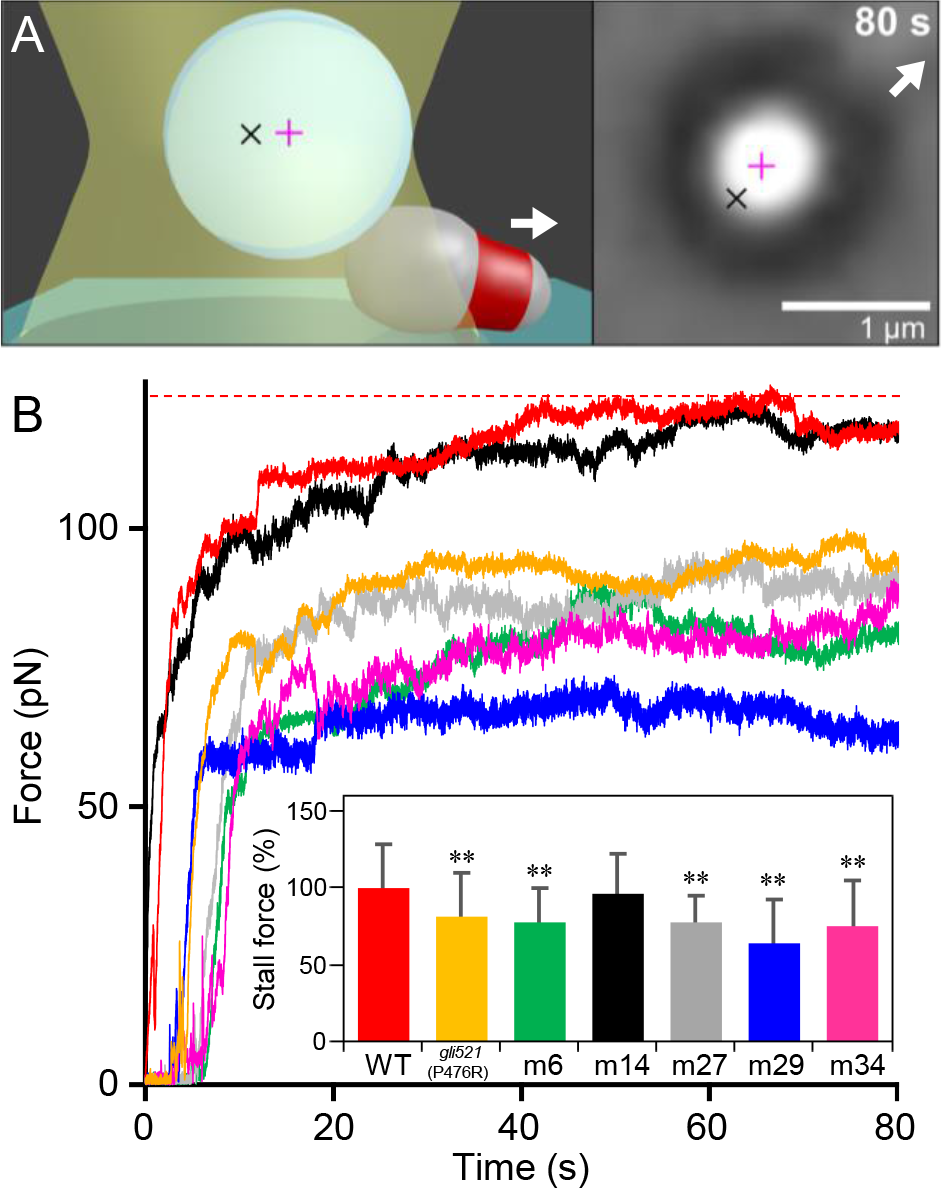
Stall force of the various strains. (*A*) Left: Illustration and micrograph of measurement using optical tweezers. The polystyrene bead (circle) (1 µm in diameter) bound to the cell is trapped by a focused laser beam (yellow hourglass) and glides in the direction of the white arrow. Black and pink crosses indicate the focal point of the laser and the bead center, respectively. Right: Optical micrographs of the trapped cell. The cell with the bead (large black ring with white center) was trapped at 0 s and stalled at 80 s. (*B*) Representative traces. The line colors correspond to the bars in the inset. Inset: Averages were normalized to the wild-type (WT) value, and are presented with standard deviations (SDs) (*n* = 50, 42, 26, 31, and 35). **, *P* < 0.01 (the difference from the WT was supported by the Student’s *t*-test). The red broken line indicates the stall force for the WT.

### Genome sequencing of various strains

To determine the proteins and mutations associated with the decreases in stall force, we sequenced the genomes of the strains-using a MiSeq sequencer. The genome of the *gli521* (P476R) mutant has been sequenced for a 30,469-bp DNA region encoding four open reading frames (ORFs): *gli123*, *gli349*, *gli521*, and *gli42*. However, the other regions remain unknown (21). The genome sequencing result for the *gli521* (P476R) mutant was consistent with the previous report of the mutation in *gli521* (21), and showed additional mutations in other regions (Table S1). One of the additional mutations causes an amino acid substitution in MvspB, a surface protein (23, 36–39). However, the reduced stall force should be caused by the mutation in Gli521, because MvspB accounts for only 1.2% of the mass of all the surface proteins, and the antibody against an abundant and closely related protein, MvspI, did not influence the gliding motility (23, 39). The genomes of the m6, m14, m27, m29, and m34 strains have not been sequenced yet, and we identified various mutations in the present study. We suggest that the decrease in the stall force in the m27 strain in the present study was caused by the mutation in *gli521*. All the strains had the same single amino acid substitution (serine to isoleucine) as the 354th residue in MMOB1700, a homolog of ABC transporter permease based on a Basic Local Alignment Search Tool (BLAST) search. This mutation may have been derived from a substitution that occurred on the clone used for the reported genome sequencing, because it was derived from the same origin (ATCC 43663) as the present study (36). Interestingly, MMOB1700 exhibited 5 other mutations in the 10 strains analyzed, suggesting a special mechanism that causes a high rate of mutation in this gene. Next, we sequenced the genomes of nonbinding strains m12, m13, and m23, which have mutations in *gli123*, *gli349*, and *gli349*, respectively (12, 15, 16, 35). Genome sequencing revealed that the identified mutations were consistent with a previous report (35), although additional mutations were identified in other regions.

### Binding and gliding of various strains

To systematically clarify the relationship between the features and mutation in the genome, we examined the binding activities and the gliding speeds for the wild-type, *gli521* (P476R), m6, m14, m27, m29, and m34 mutants, which can glide. The cell suspensions were adjusted to the same optical density and inserted into a tunnel chamber. After 15 min, we videoed the cells to count the numbers on the glass for the bound-cell ratio, and to determine their gliding speed, as previously reported (Fig. 3*A*) (29, 35, 40). The binding activities and the gliding speeds were averaged for 20 independent images and 100 cells, respectively. The binding activity and gliding speed of the *gli521* (P476R) mutant were consistent with those reported in the previous study (35). Other strains have not been analyzed by the method used here. The characteristics of the binding activities allowed classification of the strains into three groups (Fig. 3*B* left): (i) m6 had 44% of the activity of the wild-type strain; (ii) m14, m27, and m29 had 80%, 92%, and 73% of the activity, respectively; and (iii) the *gli521* (P476R) and m34 mutants had 159% and 145% of the activity of the wild-type strain, respectively. We compared these data to the binding activities previously estimated from hemadsorption, the adsorption of erythrocytes onto the surface of colonies (12). The hemadsorption values of m6, m14, m27, m29, and m34 were 24%, 93%, 113%, 96%, and 122%, respectively; i.e., except for m27, they were consistent with the results of the analyses in the present study. The gliding speed of the wild-type strain was 3.7 ± 0.2 µm/s and the relative gliding speeds of the mutants ranged from 80% to 103% of the wild-type strain, showing that the gliding speeds differed less than the binding activities (Fig. 3*B* right). Of the strains enhanced for binding, the *gli521* (P476R) and m34 mutants had reduced stall forces, suggesting that stall force is not determined simply by binding activity. Interestingly, we noticed that the proportions of the nongliding bound cells were much higher in the m6, m14, m27, and m29 strains than in the wild-type strain. The proportion of nongliding cells in the cells on glass was 6% for the wild-type strain, as previously reported (40), but those for the m6, m27, and m29 strains were higher (18%, 19%, and 16%, respectively), and that of the m14 strain was much higher (62%) (Fig. S1). This observation can be explained by assuming that gliding can be achieved by the concerted action of many molecules and interactions.

**Fig 3.**
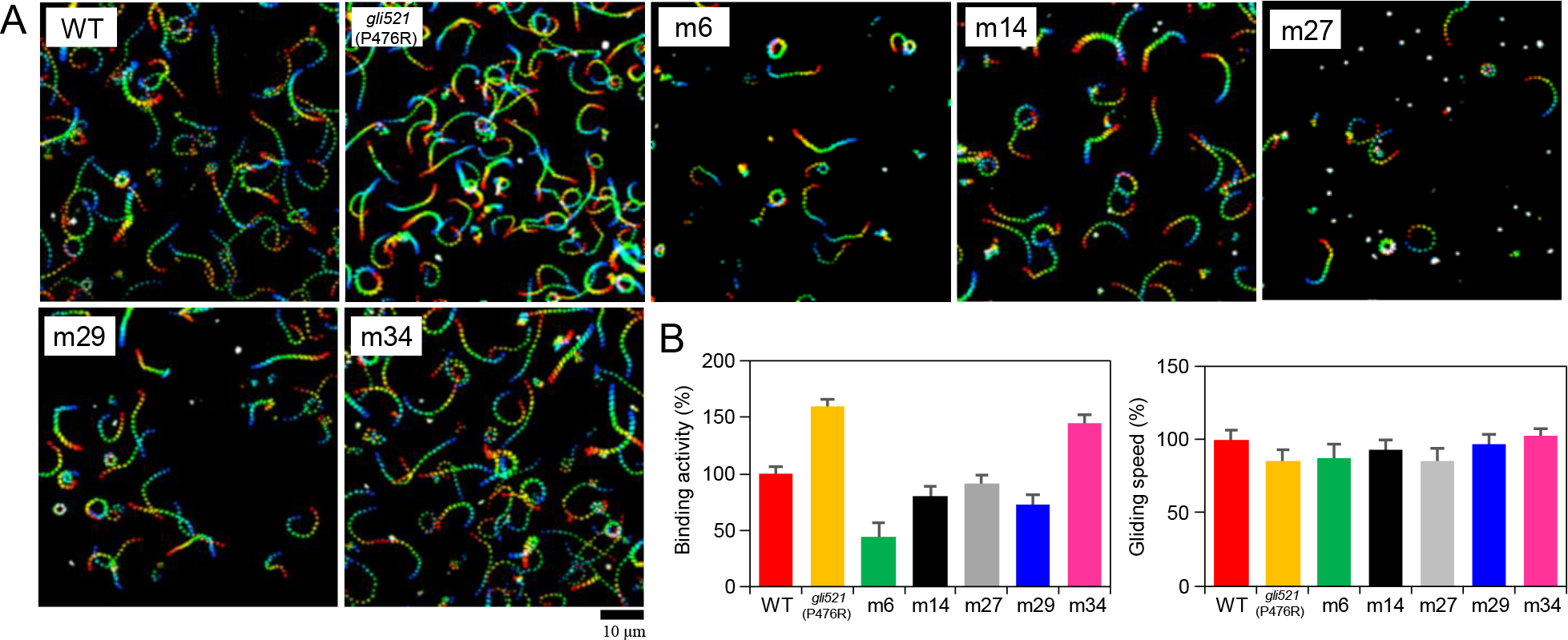
Binding and gliding properties of the various strains. (*A*) The cell trajectories are presented as a stack for 5 s, changing color from red to blue. (*B*) Averages of binding activity (right) and gliding speed (left) were normalized to the WT value and are presented with SDs.

### Effect of SOs on stall force

An *M. mobile* cell glides as a result of the integrated movement of its many legs. However, if the number of working legs is reduced, it is possible to detect the pulling force of single unit. Previous studies have shown that the number of working legs can be reduced by adding free SOs (9, 22, 25, 27, 40, 41). We therefore added various concentrations of free sialyllactose (SL)—an SO—to gliding *M. mobile* cells, and measured the stall force at 200 and 500 frames/s with 0–0.25, 0.33, and 0.5 mM free SL. The cells reached a plateau after 30–40 s, and the time taken to reach the stall increased with the concentration of free SL. The stall force decreased from 113 to 19 pN following the addition of up to 0.50 mM free SL (Fig. 4*A* and *B*). These results suggest that the stall force is the sum of the pulling force of many units.

**Fig 4.**
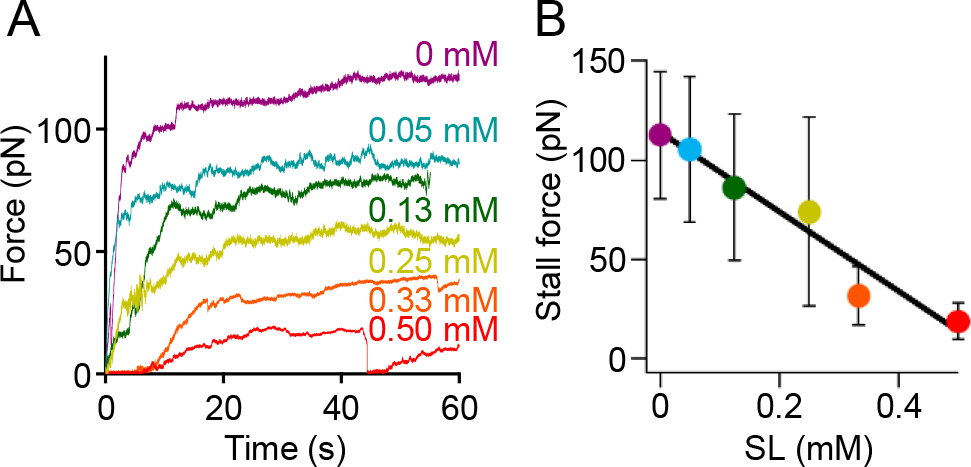
Effect of SL on stall force. (*A*) Representative traces at various concentrations of SL. (*B*) Concentration dependency on SL. Averages were plotted with SDs (*n* = 50, 13, 23, 16, 16, and 11 for 0, 0.05, 0.13, 0.25, 0.33, and 0.50 mM, respectively).

To detect the pulling force attributable to smaller numbers of units, we reduced the laser power of the optical tweezers to achieve a higher trace resolution. In this experiment, we applied 0.25 mM SL with a reduction of the trap stiffness of the optical tweezers from 0.5–0.7 to 0.1 pN/nm, whereby the trapped cells were able to escape from the trap center, and then measured the force under these conditions. The force increments with repeated small peaks were detected, and the peaks were measured using a peak-finding algorithm, as summarized in Fig. 5*A*. The distribution of these increments was determined by fitting the sum of four Gaussian curves whose peaks were positioned at 1.1, 2.0, 3.2, and 4.4 pN (Fig. 5*B*). The peaks were positioned at twice, three, and four times the value of the first peak, suggesting that these peaks reflect single, double, triple, and quadruple the minimum force increment (42). In the same way, the individual increments were analyzed for the *gli521* (P476R) mutant, and were also determined by fitting the sum of four Gaussian curves whose peaks were positioned at 0.9, 1.8, 2.6, and 3.5 pN (Fig. 5*B*).

**Fig 5.**
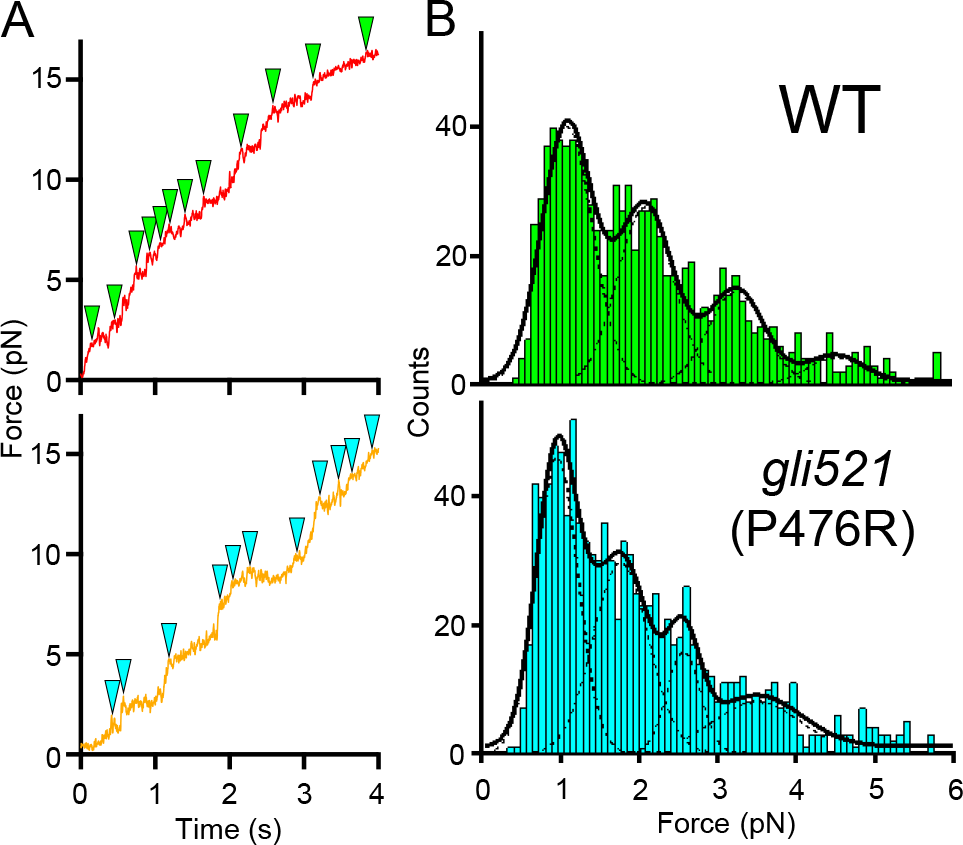
Detection of force increments under low load and 0.25 mM SL. (*A*) Representative traces of force transition in the WT and *gli521* (P476R) mutants are shown in the upper and lower panels, respectively. Green and cyan triangles in each panel indicate small peak positions taken by a peak-finding algorithm adapted to IGOR Pro 6.33J. (*B*) Distributions of peak values detected by the peak-finding algorithm were fitted to the sum of four Gaussian curves. The first, second, third, and fourth tops of the Gaussian curves were 1.1, 2.0, 3.2, and 4.4 pN in the WT, and 1.0, 1.8, 2.6, and 3.5 pN in the *gli521* (P476R) mutant, respectively (*n* = 976 and 1067).

### Stepwise force increments

Because the force increments were detected for limited leg numbers, it was possible to derive the force increments from the force generated by a single leg or a minimum force generation unit. Next, we added 0.50 mM SL to limit the number of working legs, and reduced the trap stiffness to 0.06–0.07 pN/nm to detect the minimum force increments more precisely. Very small ratios of cells remained on the glass surface under these conditions, and we attached a bead to the cells using the optical tweezers. Displacements were detected in 51 of 63 cells. Eight cells glided by creeping displacements with occasional discontinuous increments, which were mostly stepwise (Fig. 6*A*) and sometimes exhibited small peaks, as shown in Figure 5. Three cells exhibited force increments with more than six continuous steps (Fig. 6*B* Movie S2). The average value of the force increments from 46 steps or peaks in 11 cell trajectories was 1.45 ± 0.44 pN (Fig. 6*C*). These results suggest that the minimum force increment was 1–2 pN, which should be the minimum unit force for gliding.

**Fig 6.**
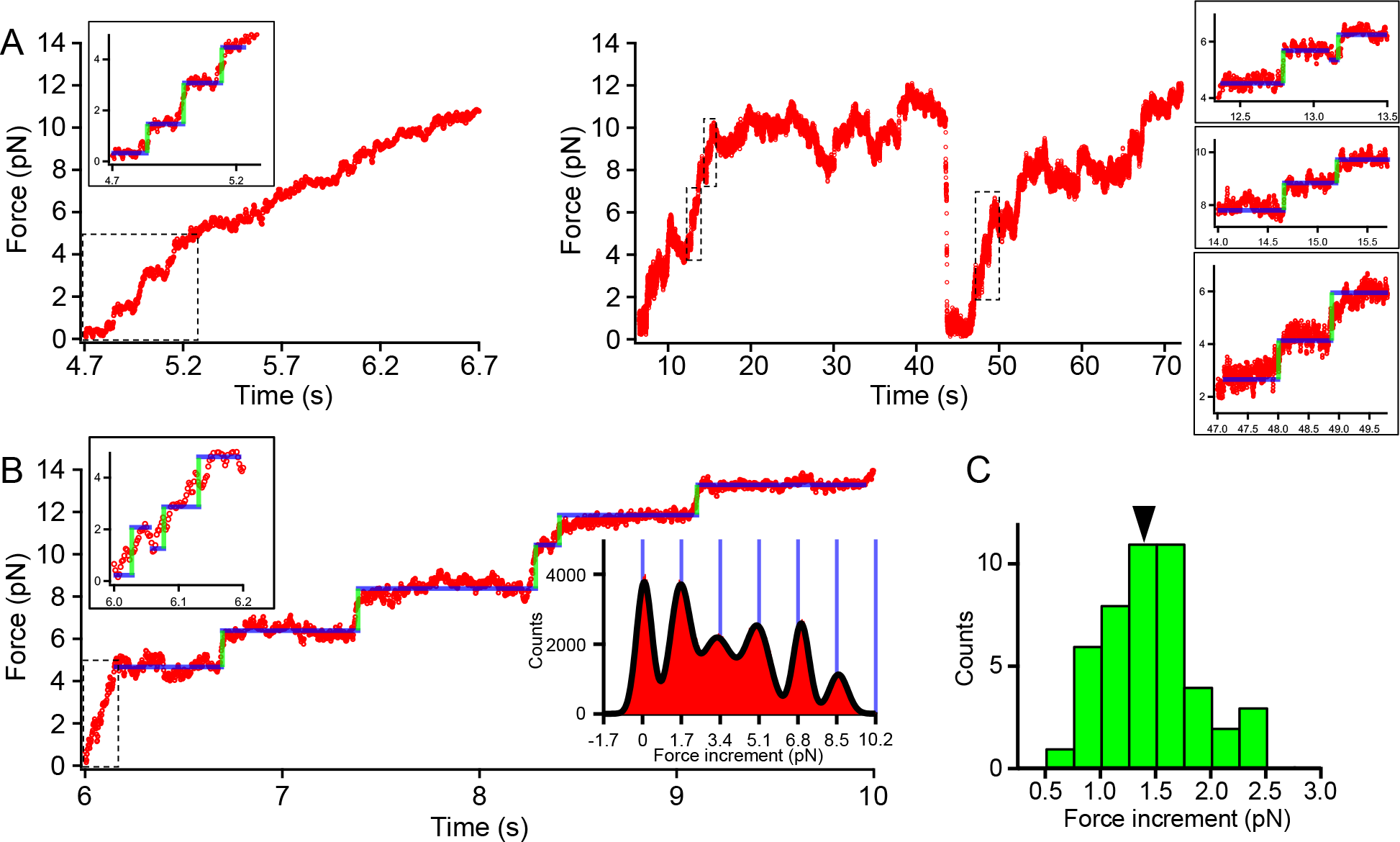
Detection of force increments under low load and 0.50 mM SL. (*A*) Two representative time courses of force generation. The trajectories in the dashed rectangular areas are magnified as insets and marked as green and blue lines for increments and dwell times. (*B*) Representative time course of continuous stepwise trajectory. The histogram of PDF analysis for indicated steps is shown in the right inset. (*C*) The histogram of force increments for 46 steps or peaks. The averaged value is indicated by a black triangle.

## DISCUSSION

### Mutations influencing gliding

In the present study, we sequenced the whole genomes of *M. mobile* mutants characterized by binding ability, gliding motility, and colony spreading. Based on these results, we propose the proteins responsible for or related to these characteristics. The m6 strain, which exhibited reduced binding, a lower gliding speed, and a weaker stall force, has mutations in FtsH, MvspI, and SecY (37–39, 43, 44). The substituted amino acid in FtsH is not conserved in other mycoplasmas except for *Mycoplasma pulmonis*, a closely related species. MvspI is probably not involved in gliding, as described above (23). SecY is generally essential for protein secretion in bacteria (44). The substitution in SecY probably affects the secretion of gliding proteins, resulting in a reduction in the binding activity of cells, because the amino acid substituted in the mutant is conserved in many mycoplasmas, including *Mycoplasma hominis*, *Mycoplasma bovis*, and *M. pulmonis*. The m14 strain was characterized by reduced binding and gliding, and its glucokinase, which phosphorylates glucose to glucose-6-phosphate at the first step of glycolysis, has an amino acid substitution (45). Interestingly, the m14 colony was less well dispersed than the wild-type colony, suggesting that glucokinase is related to gliding including chemotaxis, because colony shape is largely influenced by motility in many bacteria (12). The m27 strain, which was characterized by a small proportion of gliding cells, has a substitution at the 1461st amino acid of a total 4727 amino acids in the coded Gli521 protein, suggesting that the structure around this position is indispensable for gliding. The m34 strain was characterized by enhanced binding, and has a substitution in the conserved amino acid of the β subunit of F-type ATPase (46). The membrane potential may be related to binding activity, because F-type ATPase is responsible for the membrane potential in *M. mobile*.

### Cell behavior in stall

Each cell was stalled using optical tweezers focused on the bead bound to the back end of the cell. What events can be expected in the stall? The stall force decreased with the addition of free SL (Fig. 4). This observation suggests that the legs repeatedly catch, pull, and release SOs, even in the stalled state (Fig. 7), because the force under free SOs should increase to the stall force in the absence of SOs if the legs do not detach in stall. In our gliding model, it was suggested that the legs detach as a result of the tension caused by continuous cell displacement in gliding (14, 28, 41). The putative detachment in the stall may suggest that directed detachment occurs with much shorter displacement than expected from a 95-nm long leg structure, because the detachment also occurs in the stall. This assumption can explain the observation that the *gli521* (P476R) and m34 mutants exhibited a smaller stall force than the wild-type strain, although they had a higher ratio of cells bound to the glass (Fig. 2 and 3). The higher ratio of bound cells was probably due to the reduced force required to detach the post-stroke legs.

**Fig 7.**
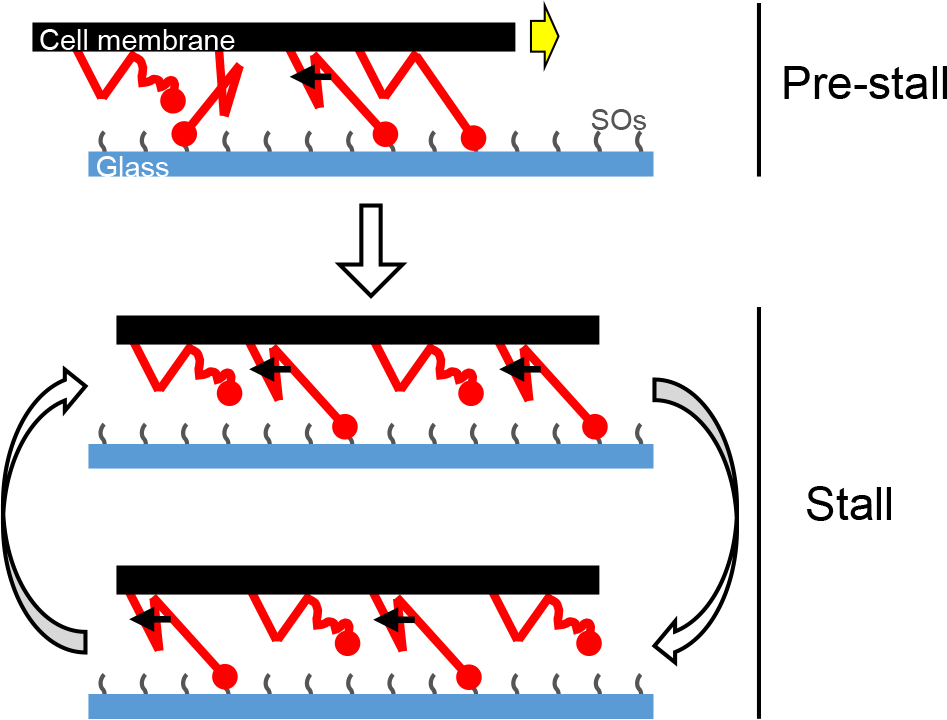
Leg behavior during stall. During pre-stall, the legs colored red repeatedly catch, pull, drag, and detach the SOs on the glass surface, thereby enabling cell propulsion. During stall, the force-transmitting legs are in equilibrium between attached and detached states. The yellow and black arrows indicate the gliding direction and the pulling force generated by the leg strokes, respectively.

### Unit number of gliding machinery

The minimum force increment did not change significantly with the distance from the laser. Because the force increments occurred additionally to the previous ones, the gliding unit should have generated the same force constantly over a rather long distance ranging from 0 to 200 nm (Fig. 6). In the present study, the stall force and the minimum force increments of a cell were approximately 113 and 1.5 pN, respectively (Fig. 2*B* and 6*C*). Previously, the number of legs in *M. mobile* was estimated to be 450 (16). The number of working units was calculated from the stall force and the minimum force increments of a cell; 113 over 1.5 pN is calculated to be approximately 75-fold, suggesting that 75 minimum units can work simultaneously. Assuming that the minimum unit of force corresponds to a single molecule of Gli349, one-sixth of the Gli349 molecules simultaneously suggested to participate in force generation in the stalled state (Fig. 1*B*). The friction occurring at the interface between an *M. mobile* cell and water flow in the interlamellar of a carp gill is calculated to be 34 pN for the maximum, based on Stokes’ law (11, 47). This number is three-fold smaller than the stall force of a cell, suggesting that a cell can glide against water flow using the force generated by the simultaneous strokes of many legs.

### *M. mobile* gliding is characterized by large steps and a small force

We compared the step size and the force of *M. mobile* gliding with those of conventional motor proteins such as myosin, dynein, and kinesin, which perform stepwise movements along the rail proteins driven by the energy derived from ATP. The step sizes and the forces of myosin-II, cytoplasmic dynein, kinesin, and myosin-V have been reported as 5.3, 8, 8, and 36 nm, and 3–5, 7–8, 8, and 2–3 pN, respectively (48–53). The long step and small force of myosin-V are caused by the lever effect of 26-nm arms (54). The step size of *M. mobile* is 70 nm; much larger than that of conventional motor proteins (22). The minimum unit force calculated here (1–2 pN) suggests the gear effect in the gliding machinery. Gli521, which is the force transmitter, forms a triskelion with 100-nm arms (24), and Gli349, which is the leg, is shaped like an eighth note in musical notation with a 50-nm flexible string (26, 55), suggesting that these proteins cause the lever effects.

### Energy conversion efficiency of *M. mobile* gliding

The direct energy source for *M. mobile* gliding is ATP; based on experiments that permeabilize cells, a “gliding ghost” can be reactivated by ATP (21). A gliding ghost exhibits stepwise movement with a dwell time that is dependent on the ATP concentration used, suggesting that the step is coupled to ATP hydrolysis (22). Based on a minimum unit force of 1–2 pN and a spring constant of 0.06–0.07 pN/nm, the work done per step, *W*_step_, is calculated to be 8– 33 pN nm from the equation *W*_step_ = 1/2 × spring constant × displacement2. Assuming that one ATP molecule is consumed per step, the energy conversion efficiency of *M. mobile* gliding can be calculated to be approximately 10–40%, because generally approximately 80 pN nm free energy is available from the hydrolysis of one ATP molecule.

F-type ATPase attains 100% energy conversion efficiency (56). It has been suggested that the gliding machinery of *M. mobile* is driven by the α-and β-subunit paralogs of F-type ATPase (18). The force transmission from this motor to the solid surfaces through several large components including Gli521 and Gli349 may be related to the putative energy loss.

## MATERIALS AND METHODS

### Strains and cultivation

The *M. mobile* strain 163K (ATCC 43663) was used as the wild-type, and its 9 mutants were grown in Aluotto medium at 25°C, as previously described (12, 35, 40, 57).

### Surface modifications of *M. mobile* cells and polystyrene beads

The cultured cells were washed with PBS/G consisting of 75 mM sodium phosphate (pH 7.3), 68 mM NaCl, and 20 mM glucose, suspended in 1.0 mM Sulfo-NHS-LC-LC-biotin (Thermo Scientific, Waltham, MA) in PBS/G, and kept for 15 min at room temperature (RT), as previously described (28, 29, 33, 34, 40). Polystyrene beads (1.0 μm in diameter) (Polysciences, Warrington, PA) were coated with avidin (Sigma-Aldrich, St. Louis, MO), as previously described (28).

### Force measurements

The avidin-coated beads were attached to biotinylated cells in two different ways, according to the concentrations of free SL used in the experiments. In force measurements under 0–0.13 mM SL conditions, the biotinylated cells were inserted into a tunnel chamber that had been precoated with 10% horse serum (20, 27, 29). Avidin-coated beads were sonicated, inserted into the tunnel chamber with various concentrations of free SL in PBS/G, and bound to the cells (32). In force measurements under 0.25–0.50 mM SL conditions, avidin-coated beads were sonicated and mixed with biotinylated cells in a microtube, and kept for 10–30 min at RT. Then, the mixture was inserted into a tunnel chamber and kept for 15 min at RT. The chamber was washed with PBS/G and the PBS/G was replaced by various concentrations of free SL in PBS/G. Both ends of the tunnel were sealed with nail polish. The bead movements were recorded at 200 or 500 frames/s and analyzed by displacement of up to 250 nm from the trap center (the linear range of the laser trap) using ImageJ 1.43u (http://rsb.info.nih.gov/ij/) and IGOR Pro 6.33J (WaveMetrics, Portland, OR) software packages (22, 28, 32, 58).

### Genome sequencing of the various strains

All the strains were plated and isolated as previously described (19). The genomic DNAs were isolated using a QIAGEN DNeasy Blood & Tissue kit (QIAGEN, Hilden, Germany). The isolated genomic DNA was sequenced using MiSeq (Illumina Inc., San Diego, CA) and mapped by CLC Genomics Workbench 8 (QIAGEN, Hilden, Germany).

### Characterization of binding and gliding of the various strains

All the strains were cultured to reach an optical density at 600 nm of 0.08. They were then suspended and inserted into a tunnel chamber (27, 29). Cell behavior was recorded and analyzed as previously reported (27, 35).

## ACKNOWLEDGMENTS

This work was supported by a Grant-in-Aid for Scientific Research in the innovative area “Harmonized Supramolecular Motility Machinery and Its Diversity” (MEXT KAKENHI; Grant Number 24117002), and by Grants-in-Aid for Scientific Research (B) and (A) (MEXT KAKENHI; Grant Numbers 24390107 and 17H01544) to MM.

---

**Table S1.**
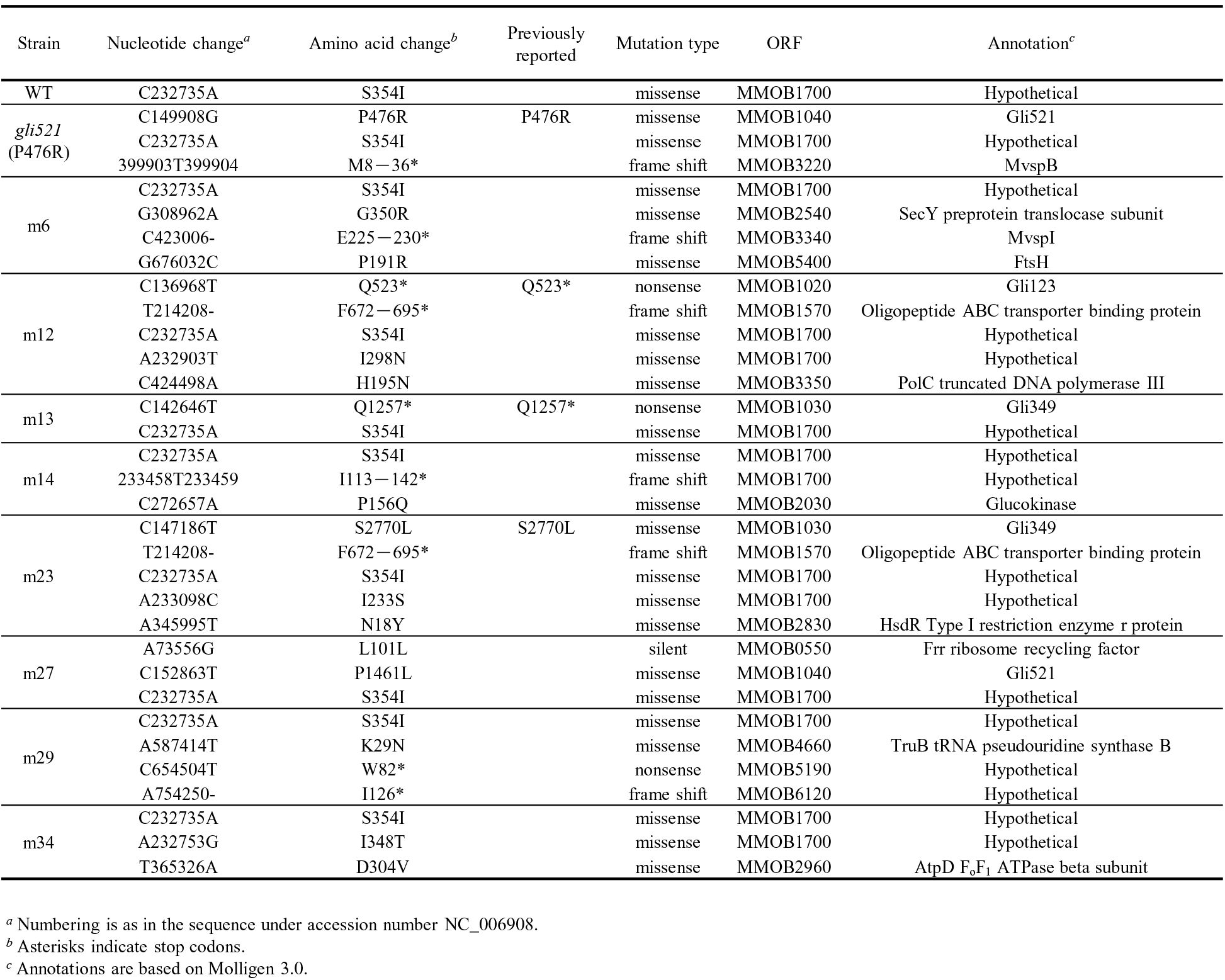
Whole genome sequence of various strains.

**Fig S1.**
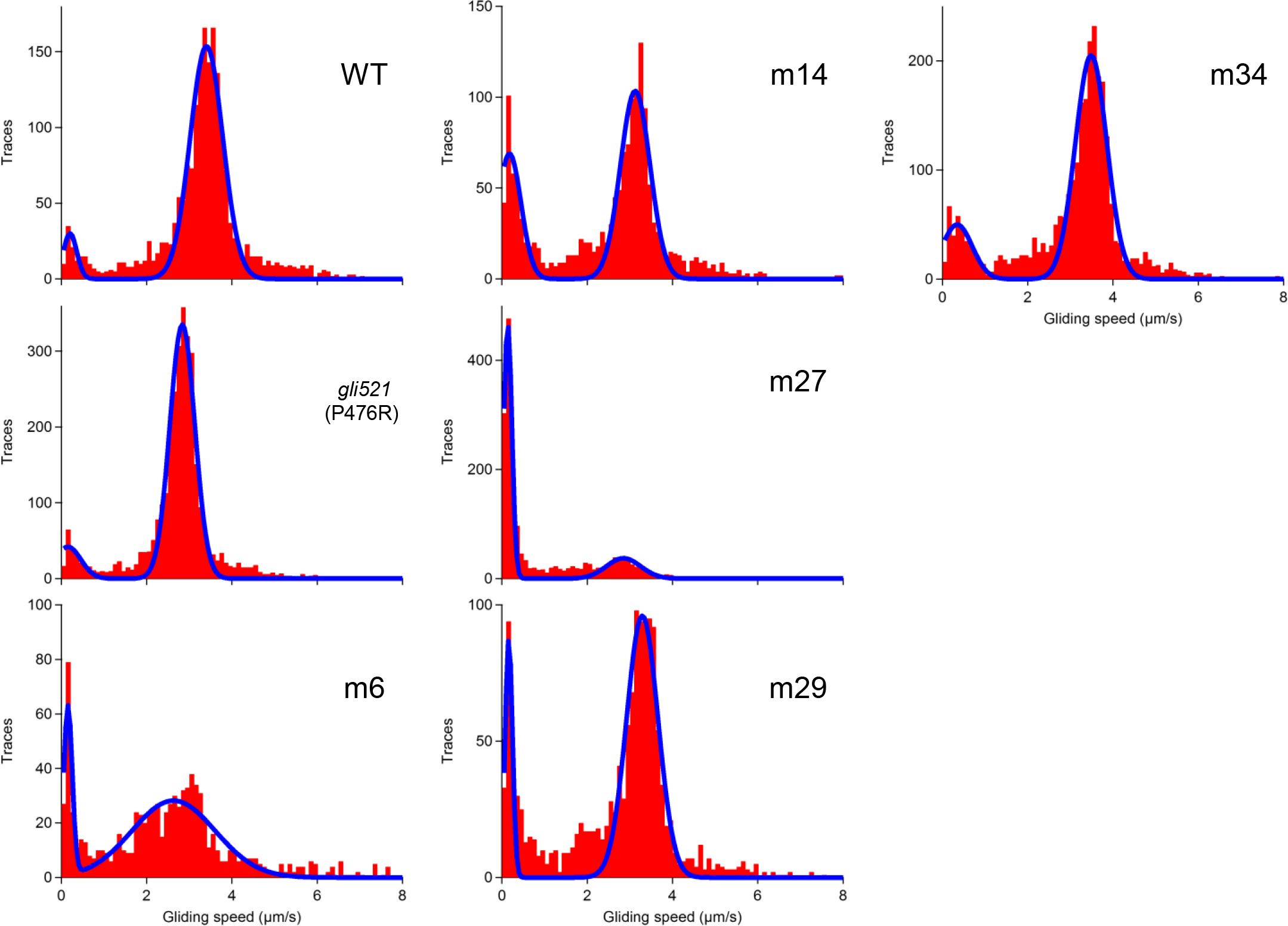
Distributions of gliding speed of the various strains over 10 s. Experimental data were fitted to the sum of two Gaussian curves colored blue.

**Movie S1.** Stall force of gliding cell carrying a bead measured using optical tweezers.

**Movie S2.** Stepwise movement of gliding cell carrying a bead detected with a weak trap.

